# Structure of Phosphorylated-like ClpXP Adaptor RssB Reveals an Interface Switch for Activation

**DOI:** 10.1101/2022.12.15.520641

**Authors:** Christiane Brugger, Jacob Schwartz, Martin Filipovski, Alexandra M. Deaconescu

**Author notes:** Address correspondence to: Alexandra M. Deaconescu, B.E., Ph.D., Assistant Professor of Molecular Biology, Cell Biology and Biochemistry, Laboratories of Molecular Medicine, Brown University, 70 Ship St. G-E4, Providence, RI 02903, USA.

## Abstract

**SUMMARY:** In γ-proteobacteria such as *Escherichia coli*, the general stress response is mediated by σ^s^, the stationary phase dissociable promoter specificity subunit of RNA polymerase. σ^s^ is degraded by ClpXP during active growth in a process dependent on the RssB adaptor, which acts catalytically and is thought to be stimulated by phosphorylation of a conserved aspartate in its N-terminal receiver domain. Here we present the crystal structure of full-length RssB bound to a beryllofluoride phosphomimic. Compared to the inhibited IraD anti-adaptor-bound RssB structure, our study reveals movements and coil-to-helix transitions in the C-terminal region of the RssB receiver domain and in the inter-domain segmented helical linker, accompanied by packing of the C-terminal effector domain onto the α4-β5-α5 (4-5-5) “signaling” face of the RssB receiver domain. This face is often the locus of protein-protein interactions in unphosphorylated receiver domains, but its masking is unusual in their phosphorylated forms. Our structure emphasizes the remarkable plasticity that underpins regulatory strategies within the large family of response regulators.

## INTRODUCTION

The ability to adapt to a changing and sometimes adverse environment is a fundamental property of any living system. It ensures survival. In bacteria, deregulation of the pathways associated with stress responses are implicated in growth defects^1^, biofilm formation^1-3^, antibiotic resistance^1,4,5^ and altered ability to colonize hosts^6,7^. By definition, general stress responses (recently reviewed by Gottesman)^1^, unlike specific stress responses, are mediated by global regulators and confer broad cross-resistance to stress signals that did not originally induce the response. In γ-proteobacteria, the general stress response is mediated by σ^s^, the stationary phase dissociable promoter specificity subunit of RNA polymerase. This turns on a large regulon of up to 24% of the *E. coli* genes^8-11^ and is widely dubbed as the master regulator of the general stress response. While increased σ^s^ levels are required for bacterial virulence and responses to stress, including antibiotic stress^12-14^, unregulated overexpression of σ^s^ can also be detrimental^15^, likely because under these conditions, the system cannot restart growth even after the stress signal has gone away. This underscores that for a plastic stress response and niche adaptation, cells must regulate intracellular σ^s^ levels in a tight and fast manner. This is accomplished through both positive and negative regulation at multiple levels, including (1) transcription activation^16-19^, (2) translation activation/repression, involving structural rearrangements of the *rpoS* mRNA by small regulatory RNAs and RNA chaperones^19-21^, and (3) protein turnover by the ATP-dependent ClpXP machinery^19,22-26^.

By itself, σ^s^ is a poor substrate for ClpXP, and needs to be presented to the protease by an adaptor, the two-domain response regulator RssB (aka MviA, SprE)^22,25-27^. Substrate presentation represents the rate-limiting step in σ^s^ turnover^28^ and is regulated by phosphorylation of D58 in the N-terminal receiver domain (RssB^NTD^)^29^. As a response regulator, RssB is atypical – its effector domain (RssB^CTD^) is specialized for protein-protein interactions^30^ rather than DNA binding as in most response regulators^31^. Moreover, while other adaptors operate in pairs with cognate anti-adaptors, RssB is unique in that it is inhibited by multiple, unrelated and stress-specific anti-adaptors (IraD, IraM, IraP and IraL)^32,33^, suggesting a great potential for plasticity and specialization of the response to distinct triggers via negative regulation.

The role of RssB phosphorylation in adaptation to stress or in the recovery from stress has remained controversial for several reasons. The phosphodonor acetyl phosphate (AcP) was found to stimulate σ^s^ degradation^27,29,32,34^, an effect that was attributed to phosphorylation promoting the formation of a stable binary σ^s^-RssB and a ternary σ^s^-RssB-ClpX delivery complex^27^. As such, *in vitro* reconstitution of σ^s^ proteolysis demonstrated that phosphorylation stimulates the degradation reaction^27,32,34^, with a ∼ threefold decrease in the apparent K_M_^30^. A recent study suggested instead that phosphorylation lowers the affinity of the RssB-σ^s^ interaction^35^, but in this case RssB and σ^s^ truncations were employed for analysis, which according to our work, do not faithfully recapitulate the interactions between the full-length proteins. The situation is further complicated by the finding that a strain carrying a non-phosphorylatable substitution of D58 on the chromosome displays no strong phenotype under either starvation or supplementation with the limiting nutrient (recovery from stress)^36^. Thus, after decades of work on this system, only very limited insights into how phosphorylation affects communication between the two RssB domains for σ^s^ binding and hand-off have been achieved.

Here we sought to clarify the effects of phosphorylation on RssB using a combination of X-ray crystallography of phosphorylated-like full-length RssB, and functional assays. We show that the molecular plasticity of RssB is, at least partially, phosphorylation-dependent and affects primarily the C-terminal α5 helix of the receiver domain and the interdomain segmented helical linker (henceforth referred to as SHL). Structural remodeling of these elements results in alternative packing of the RssB constituent domains. While activation of response regulators often involves disruption of inhibitory interactions between the receiver and effector domains to liberate their 4-5-5 face for target binding^37^, we unexpectedly observe that RssB phosphorylation induces a switch in the interface used for effector domain packing, from the α1− α5 face in the non-phosphorylated inhibited state (i.e. bound to IraD) to the 4-5-5 face in the phosphorylated-like free state. Thus, the 4-5-5 face, the canonical locus for receiver domain oligomerization and association with targets, is utilized for an interface switch to achieve an alternative closed RssB structure, which can nevertheless associate with σ^s^ with high affinity and may play a role in substrate hand-off to ClpXP.

## RESULTS

### Trapping Phosphorylated-like RssB with a Phosphomimic

Despite significant efforts, full-length RssB has long resisted crystallization due to its limited solubility, and what is thought to be a dynamic nature, characterized, as in other response regulators, by a dynamic equilibrium of “OFF” forms^38^ – non-phosphorylated and with poor affinity for σ^s^ - and an “ON” form – stabilized by phosphorylation and with high affinity for σ^s^. Examples in which the active or partially active form of response regulators can be accessed spontaneously by conformational breathing, in the absence of phosphorylation have also been reported^38-40^. In the case of RssB, most of our understanding of phosphorylation-dependent activation comes from studies employing AcP to stimulate formation of an σ^s^:RssB:ClpX assembly^27^ and promote σ^s^ degradation without being an absolute requirement for it^27,32,34,41^. We first used our high-quality preparations of pure RssB to verify RssB activity by performing *in vitro* phosphorylation assays coupled to Phos-tag electrophoresis. In our phosphorylation assays (Figure 1A), RssB was rapidly modified within the first 5 minutes of incubation with AcP, but 100% phosphorylation was not achieved even after 40 minutes of reaction time, likely due to the spontaneous autodephosphorylation activity of RssB^NTD^, a general property of receiver domains, which can vary by orders of magnitude from protein to protein^42^. Consistent with earlier work^27^, we found that in the presence of AcP and Mg^2+^, RssB forms a 1:1 stable complex with σ^s^ even in the absence of ClpX (Figure 1B). No binary RssB: σ^s^ or ternary σ^s^:RssB:ClpX complexes were observed in the absence of AcP, as reported before^27^, suggesting that dephosphorylation may play a role in substrate hand-off and/or RssB recycling. We carried out extensive crystallization trials in the presence of AcP, but these remained unfruitful. Because the phosphoaspartyl bond is labile in aqueous solutions and receiver domains have intrinsic dephosphorylation activity^43^, we next sought to limit sample heterogeneity by replacing AcP with Mg^2+^•beryllofluoride. This phosphomimic has been used for trapping phosphorylated-like conformations of numerous response regulators^44^, including the truncated receiver domain of RssB, RssB^NTD 45^. As expected, beryllofluoride addition led to a substantial increase in the amount of of RssB pulled down by His-tagged σ^s^ that was readily apparent in a ∼5-fold increase in pull-down efficiency in the presence of Mg^2+^•beryllofluoride rather than in its absence (Figure 1C, compare lane 3, left to lane 3, right).

**Figure 1.**
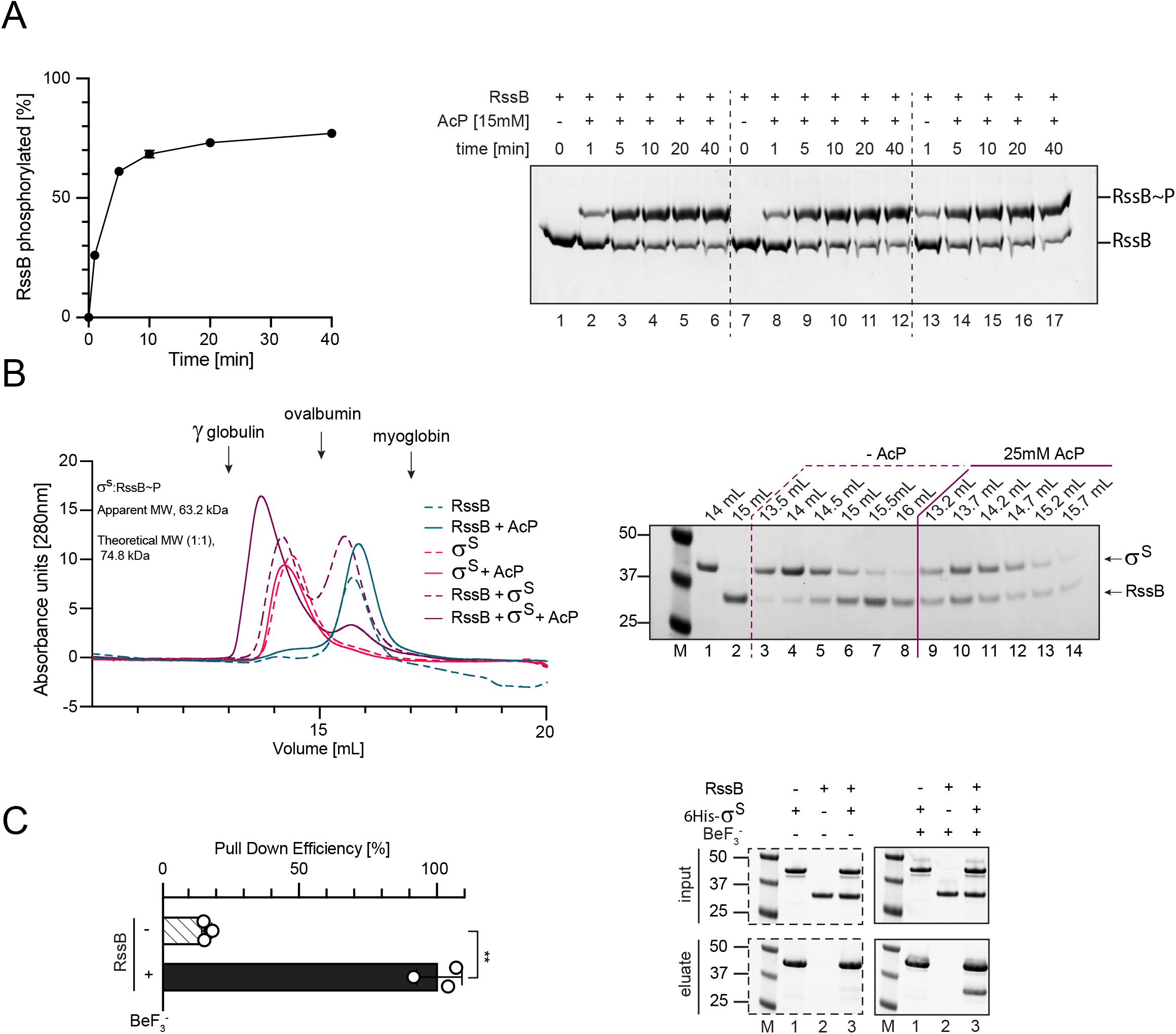
Trapping RssB∼P with a Phosphomimic. A. Time course of RssB phosphorylation. Reactions were initiated by addition of 15 mM AcP, mixed at various timepoints with LDS-loading buffer and loaded onto 10% Phos-tag gels (FujiFilm Wako). Shown are means ± s.d. (left, n=3) together with one representative Phos-tag gel containing three replicates (right). Symbols are generally smaller than error bars. B. Size-exclusion chromatography profiles of RssB, σ^s^ and premixed RssB: σ^s^ complexes ± 25 mM AcP and 10 mM Mg^2+^. Insets show SDS-PAGE analysis of peak fractions eluted off a Superdex 200 10/300 column. Phosphorylation was initiated by pre-incubation with AcP for 15 min on ice prior to injection. C. Pull-down assay probing the RssB-σ^s^ interaction in the absence and presence of beryllofluoride and AcP. Purified RssB was pulled down via 6His-σ^s^, which was immobilized to Ni^2+^-NTA beads. Shown are means ± s.d. (n greater or equal to 2) together with one representative SDS-PAGE analysis. Statistical analysis was performed using a two-tailed Student’s t-test (**, p<0.01).

To limit aggregation of RssB, we performed crystallization screening at protein concentrations regimes well below standard ones (Materials and Methods) and obtained well-diffracting crystals of phosphorylated-like RssB, RssB∼P (Table 1), but not RssB. Phasing was achieved using molecular replacement with individual domains derived from the structure of the IraD-bound RssB^D58P^ (PDB ID 6OD1)^34^.

**TABLE 1.**
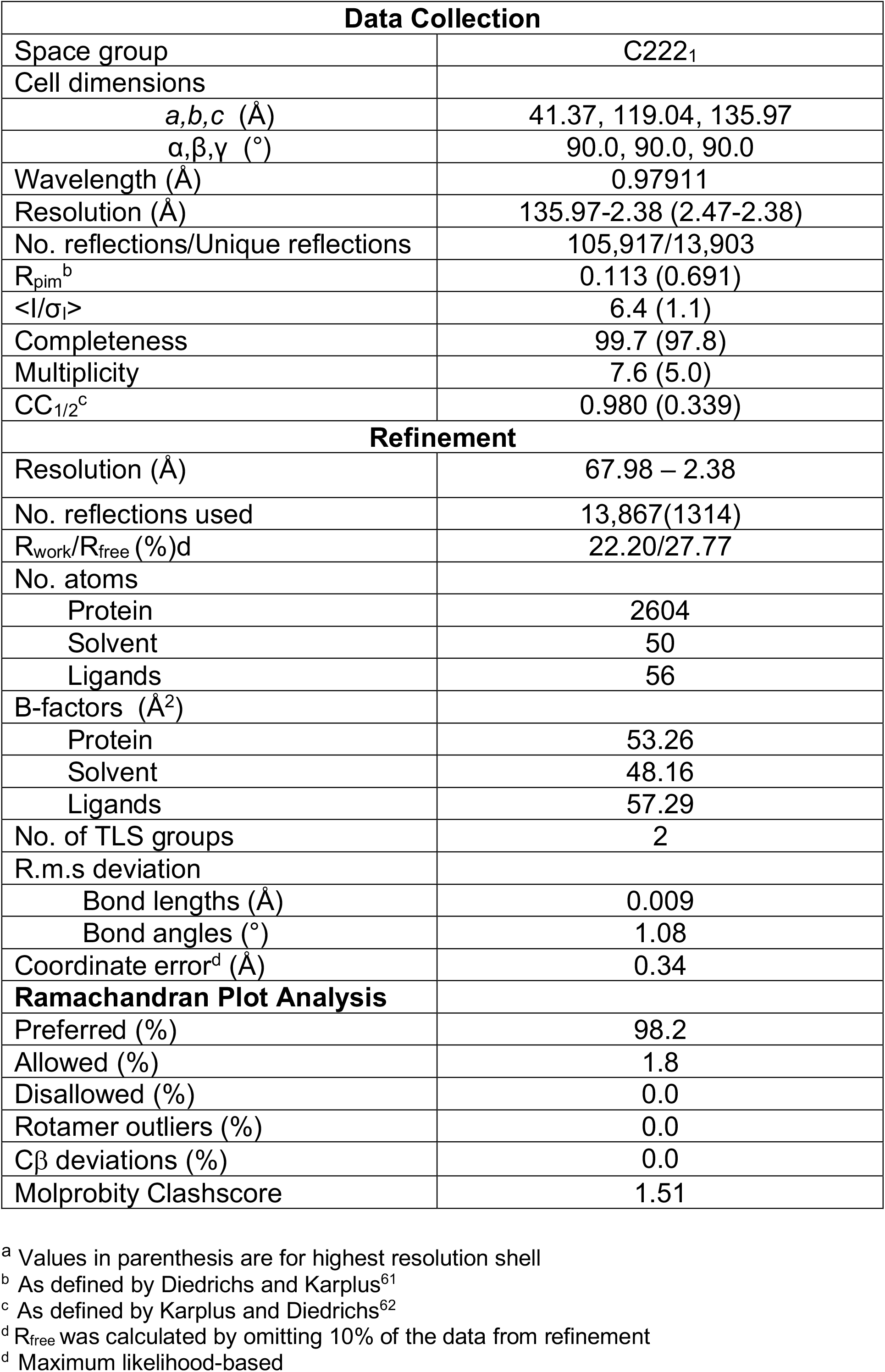
X-ray Data Collection and Refinement Statistics^a^.

### Architecture of Beryllofluoride-bound RssB

Beryllofluoride-stabilized RssB assumes a compact conformation with contacts between RssB^NTD^ and its C-terminal pseudophosphatase (RssB^CTD^) domain that bury an interface of approximately ∼529 Å^2^, suggesting a transient and low-affinity interaction between the two domains. The “closed” conformation observed here is stabilized by the SHL (magenta in Figure 2) that docks atop of the proximal edge of the 4-5-5 face and packs against α8, holding the two domains together (Figures 2B, C). The Mg^2+^-beryllofluoride moiety is well ordered and surrounded by well-conserved residues forming a quintet required for phosphorylation (dotted outline cartouches in Figure 2A and sticks in Figure 2D) and well-ordered water molecules, which complete the octahedral coordination of the Mg^2+^ ion.

**Figure 2.**
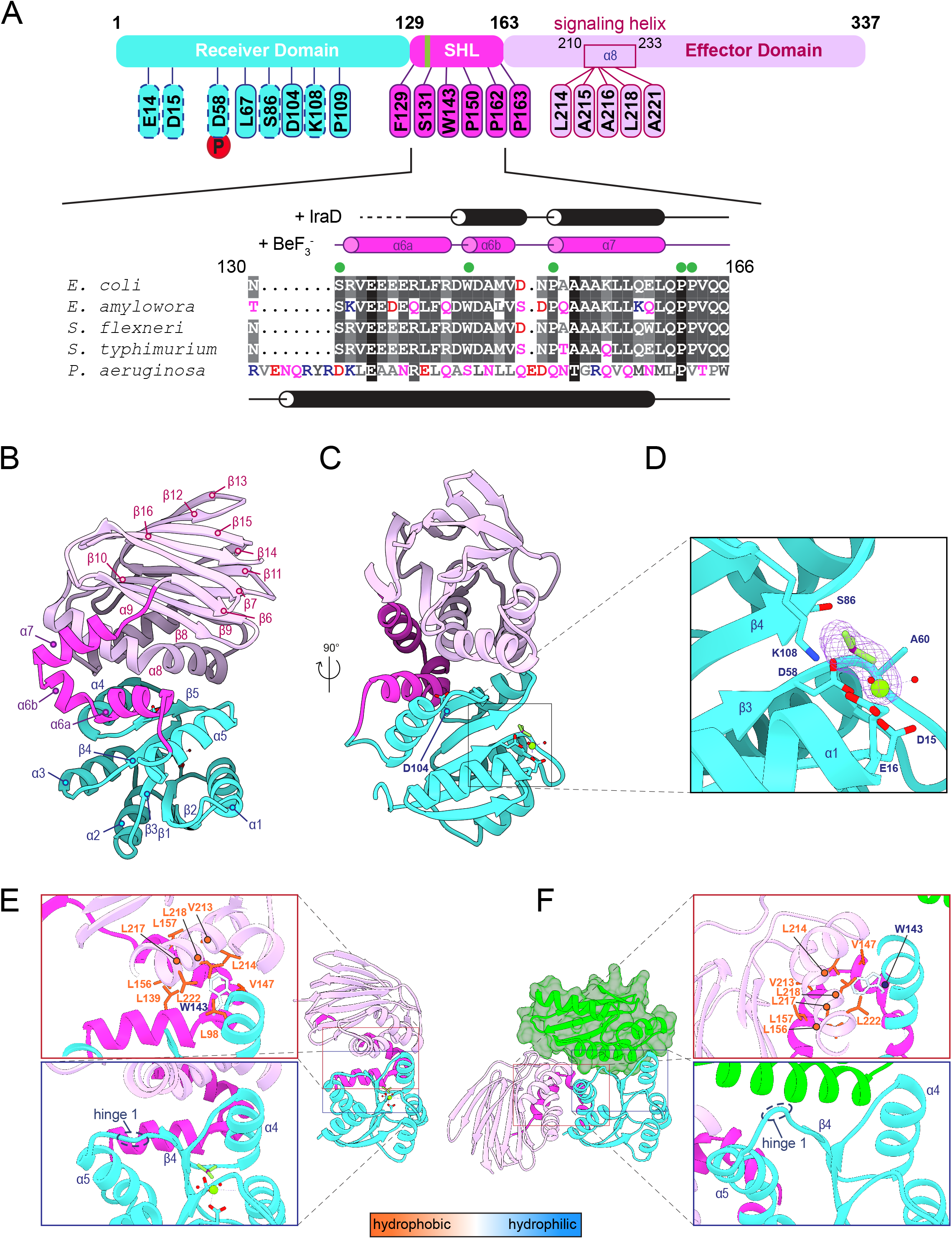
Overall architecture of RssB·BeF_3_ ^-^•Mg^2+^. A. Domain architecture of *E. coli* RssB. Phosphoacceptor D58 is indicated with a red circle and key functional residues are highlighted as cartouches. Residues belonging to the phosphorylation stabilization quintet are marked by red outline cartouches. The polyglutamate motif within the SHL (magenta) is shown as a green vertical bar. Selected SHL sequences shown below are colored by evolutionary conservation from white (not conserved) to black (invariant). Secondary structural elements in the presence of BeF_**3**_ ^-^ and IraD, as defined with DSSP, are also shown together with the locations of hinges 3-6 in SHL (green circles). Hinges 1-2 in RssB^NTD^ are not shown here. The SHL is well conserved among species in which RpoS proteolysis has been experimentally demonstrated^22,25,26,57-59^, but not between *E. coli* and *P. aeruginosa*. These proteins appear to function differently^32,34^. B-D. Top (B) and side (C) views of RssB•BeF_**3**_ ^-^•Mg^2+^ colored by domain as in A. The phosphoacceptor residue, D58, is shown as sticks and magnified, together with the beryllofluoride moiety in panel D. Density in the Polder omit map corresponding to the beryllofluoride ligand is contoured at σ and the signaling quintet residues (D15, E16, D58, S86, K108) that coordinate the Mg^2+^-beryllofluoride moiety and are required for phosphorylation-dependent signaling are also shown as sticks. Water molecules coordinating Mg^2+^ (green) are shown as red spheres. Beryllofluoride is shown in stick representation. E,F. Side-view of beryllofluoride-bound RssB (E) and IraD-bound RssB^D58P^ (F, PDB ID 6OD1) colored by domain organization as in A. Residues shows as sticks are colored according to hydrophobicity using the Kyte-Doolittle scale^60^. Structures were oriented based on a superposition restricted to the Cα–trace of RssB^NTD^ (r.m.s.d of 0.840 Å). The proteolytically stable IraD truncation, IraD^Trunc^, docks primarily on the 4-5-5 face of RssB^NTD^ and is shown as a green transparent surface in F. Insets highlight the conformational differences in α– and the hinge 1 region around K108 and P109. RssB^NTD^/RssB^CTD^ and SHL interfaces in E are predominantly hydrophobic as indicated by the clustering of hydrophobic residues (Ile, Val, Leu). Among these are well characterized residues (e.g. L214, L218 and L222), which when mutated lead to resistance to anti-adaptor regulation, but not anti-adapter binding and hyperactivate RssB, possibly due to enhanced interactions with σ^s 32^. Also shown is W143, whose sidechain inserts itself, in both structures, in a cage of hydrophobic residues, including L214 and L218.

Interdomain interactions as well as SHL-domain interactions are driven mainly by van der Waals contacts, with few interfacial hydrogen bonds observed at our resolution and many highly conserved hydrophobic residues on the interior faces of the central helical bundle at the RssB^NTD^/RssB^CTD^ interface (Figure 2E), suggesting a stable hydrophobic core. Some of these hydrophobic interactions map the helix α8, where Battesti et al. identified a series of residues (e.g. L214, A216, L218) whose substitution results in resistance to inhibition by anti-adaptors and may mimic the effect of phosphorylation^32^. These are also seen to be part of a hydrophobic “cage” that surrounds W143 (Figure 2E, F), a residue with hinge-like potential.

The RssB structure observed here differs from the “closed” structure of RssB^D58P^ bound to IraD (Figures 2F)^34^. Pairwise superposition via the Cα–traces of the RssB^NTD^ domains reveal large structural deviations that start with the loop preceding helix α5, around residues K108/P109 (Figure 2E, F), and become amplified along the SHL due to the swiveling motion of the SHL segments that result in a 42 Å translation and 57º rotation of RssB^CTD^ relative to RssB^CTD^ in the IraD-RssB^D58P^ complex (Figures 2E, F; 3B, C and Movie 1).

### Beryllofluoride Binding Does Not “Unpack” RssB into an Open Structure with Well-Separated Domains

We have previously proposed that the SHL is plastic and serves as the primary target for both positive (phosphorylation-mediated) and negative (anti-adaptor mediated) regulation of RssB, which becomes stabilized in either an “open” or “closed” state, respectively^34^. Consistent with high structural plasticity, we observe several differences in the conformation of α5 and the SHL relative to the IraD-bound RssB^D58P^ structure. These changes are organized around five potential hinge regions (Figure 3A) that articulate the segments of the SHL and which are highly conserved among RssB orthologs experimentally confirmed to mediate RpoS proteolysis (Figure 2A), and not conserved in the *Pseudomonas aeruginosa* RssB-like protein. The latter represents the closest RssB homolog of known structure (PDB IDs 3EQ2 and 3F7A, unpublished), and adopts an open dumbbell architecture stabilized via dimerization mediated by a central coiled coil with characteristic knobs-into-holes interactions that are absent in RssB, as noted by Dorich et al.^34^.

**Figure 3.**
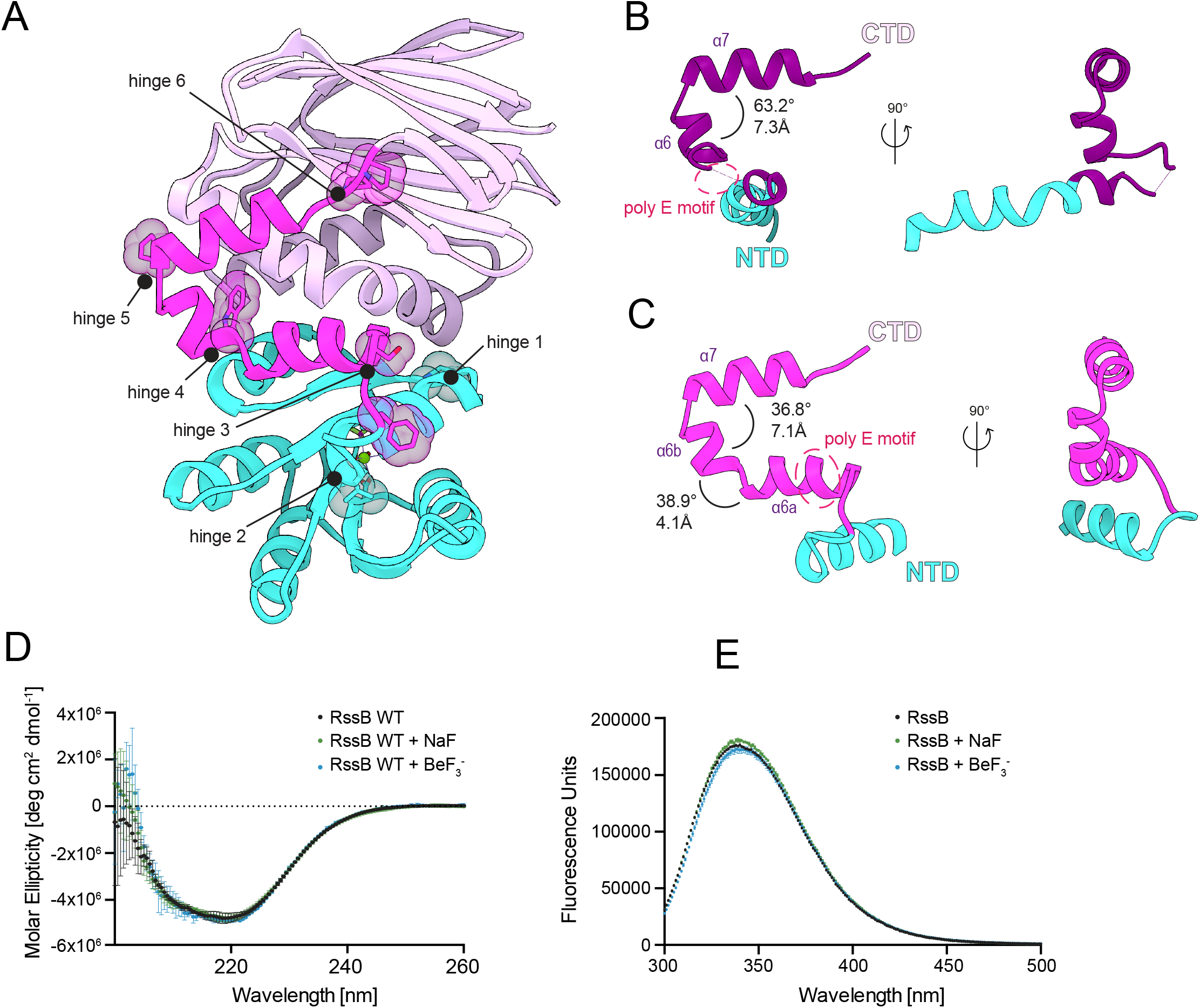
RssB is organized around 6 hinges and features an SHL that undergoes disorder-to-order transitions. A. View of RssB with residues that serve as potential hinges highlighted in CPK representation. B,C Side-by-side view of the SHL in the IraD-RssB^D58P^ complex (dark purple, B) and beryllofluoride-bound RssB (magenta, C). Structures were oriented based on a superposition of RssB^CTD^ (not shown for simplicity), which results in a good alignment with α7, but not of α–, α6a and α6b. D. Circular dichroism spectra of RssB at 2.5μM in presence/absence of 1mM beryllofluoride and 6mM NaF. Spectra were acquired as described in Materials & Methods. Error bars represent s.d. (n=3 technical replicates). E. Intrinsic Trp fluorescence spectra collected on 2μM RssB in the presence/absence of 1mM beryllofluoride and 6mM NaF. Spectra acquisition is described in Materials & Methods. Error bars represent s.d. (n=3 technical replicates).

First, hinges 1 and 2 (P109 and L124) demarcate α5, which is shorter, straight and displaced, contrasting the kinked conformation seen when bound to IraD (Figure 3 and Movie 1). A kinked-to-straight conformational transition likely sterically impinges on the RssB^CTD^ upon phosphorylation, may force it to repack and weaken IraD contacts. While IraD can bind both RssB and RssB∼P, its affinity for RssB∼P is lower^35^, in agreement with the structural differences and RssB^CTD^ docking onto the 4-5-5 face, for which IraD must compete for binding^34^.

The last four hinges are located either in the loop preceding SHL (S129; F131) or along the SHL (Figure 2A and 3A). The SHL largely maintains its helical segmented structure, with the C-terminal α7 helix strongly coupled to RssB^CTD^, together with which it moves as a rigid body relative to RssB^NTD^. At its N-terminus, however, the linker undergoes a disorder-to-order transition such that its polyglutamate motif (residues E134-E137) folds, and adopts α–helical psi/phi combinations. The rest of the hinges are positioned where α6 splits into α6a and α6b (W143, hinge 4), in between helical segments α6b and α7 (P150, hinge 5) or on the loop connecting to RssB^CTD^ (P162;P163; hinge 6). The importance of some of these hinge residues was previously assessed by Battesti et al., who determined that variants carrying substitutions of W143 and P150 are competent for adaptor activity *in vivo* and *in vitro*, while variants carrying substitutions in the N-terminal SHL region (RssB^P109S^) are slightly defective^32^. Perhaps most striking is the pronounced functional defect caused by substitution of K108^30,32^, a highly conserved residue critical for general signaling by receiver domains^46^. The mechanistic underpinnings of the defects seen in the K108 variants remain unknown, but since phosphorylation is not an absolute requirement for σ^s^ degradation^24,27,34,36,41^, it appears likely that K108 might play roles in addition to signaling, such as in protein-protein interactions with either σ^s^ or ClpX. Substitutions in the polyglutamate motif (E135A; E136A; E137A, resulting in the RssB^AAA^ variant) are also detrimental to the adaptor function of RssB both *in vitro* and *in vivo*, yet they do not greatly affect stimulation of degradation by phosphorylation^34^, suggesting that phosphorylation-induced conformational switching can still be achieved with this variant and that perhaps the polyglutamate motif may also contribute to protein-protein interactions. A similar defect in adaptor function was also observed for a variant carrying an alanine substitution of the 4-5-5 face residue D104, seen in Figure 2C.

To further probe for large-scale, global conformational changes that might be induced by phosphorylation, we employed circular dichroism (Figure 3D) and intrinsic tryptophan fluorescence spectroscopy (Figure 3E) in the absence/presence of beryllofluoride. None of these experiments indicated beryllofluoride-dependent changes that would be expected upon “opening” of the molecule. Previous analysis using limited proteolysis also failed to indicate differences in susceptibility to protease treatment in the presence/absence of AcP^34^. We therefore conclude that phosphorylation does not induce large global conformational changes that would stably alter the packing of the two domains, at least in the absence of σ^s^, but rather local changes, mostly in the receiver domain and perhaps the SHL.

## DISCUSSION

RssB, the sole adaptor responsible for delivery of σ^s^ to the ClpXP protease was identified more than two decades ago as the first response regulator to function as an adaptor and mediate protein-protein interactions with ClpX^22,26,27^. Shortly thereafter, it was shown that D58 within its receiver domain is the sole phosphoacceptor site^29^. Despite multiple attempts to identify the RssB kinome, the source of *in vivo* RssB phosphorylation has remained mysterious. Kinase ArcB and small phosphodonor AcP have been implicated ^47,29^, but it appears likely that RssB may be phosphorylated by multiple yet-to-be-defined kinases, possibly ensuring redundancy and efficient destruction of σ^s^ during active growth or upon return to active growth. Phosphatase activities of bifunctional kinases may also be involved in shifting the equilibrium towards the OFF form, particularly under conditions when σ^s^ levels must increase (Figure 4). Chromosomal substitution of the phosphoacceptor aspartate with non-phosphorylatable alanine, while increasing the σ^s^ half-life fivefold, did not result in a strong phenotype under selected conditions of either starvation or recovery from stress^36^. This is somewhat surprising given that under active growth, RssB levels are low (on the order of 3-4 molecules per average cell cycle and barely detectable by immunoblotting^48^), so one would expect that RssB must have a high affinity for its target to effect any change, which is not the case in the absence of phosphorylation (Figure 1A). However, the RssB promoter is σ^s^-controlled and under homeostatic control due to σ^s^ degradation^28^, and so because σ^s^ levels fluctuate, it is possible that phosphorylation is critical only under specific conditions and during a narrow time window. Interpretation of previous work is also complicated by the use of multi-copy plasmids, which can alter the ratio of RssB: σ^s^ from what is typically encountered physiologically^49^.

**Figure 4.**
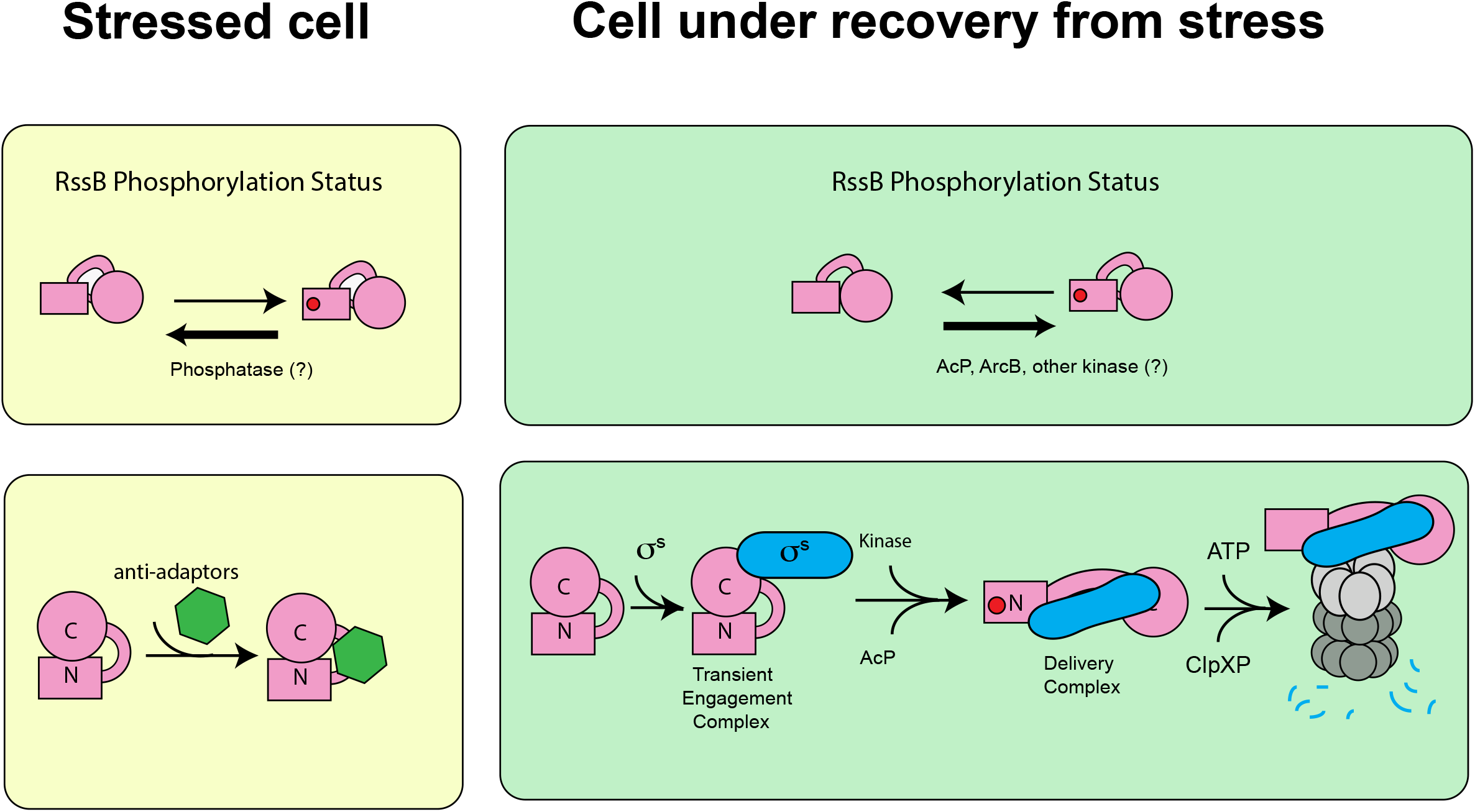
Molecular Events Leading to RssB Activation and σ^s^ Degradation. Following the two-state equilibrium model generally applicable to response regulators, we propose that RssB assumes OFF conformations (with low affinity for σ^s^) and an ON conformation (with high affinity for σ^s^), which is promoted by phosphorylation of D58 (red circle). Conformational differences between the ON/OFF forms may be subtle and are not explicitly indicated here. Upon stress, RssB is inhibited by anti-adaptors, which prevent σ^s^ proteolysis. Upon recovery from stress, we propose that RssB engages sequentially with σ^s^ via the C-terminal domain first in a transient engagement complex, for which phosphorylation is not a requirement, and then via the N-terminal domain via a delivery complex, the formation of which is promoted by phosphorylation. Upon delivery to ClpXP, hydrolysis of the acyl phosphate bond signals hand-off of the substrate to ClpXP, and subsequent recycling of RssB.

Our study circumvents some of the limitations of previous studies by trapping wild-type full-length RssB with the beryllofluoride phosphoryl mimic and by revealing changes in protein plasticity (Figures 2-3). While other models cannot be ruled out in the absence of a structural model for free, unphosphorylated RssB, our data are suggestive of phosphorylation modulating the RssB structure locally, particularly in helix α5 and the SHL. Then, how does phosphorylation stimulate degradation?

While speculative at this point, we favor the following sequence of events. Phosphorylation leads to conformational changes around D58, which preorganizes a high affinity binding site for the σ^s^. We propose that σ^s^ binds initially to RssB^CTD^ and possibly the SHL regardless of whether RssB is phosphorylated or not, in agreement with previous findings^30^. Upon docking of σ^s^ to RssB^CTD^ (“transient engagement complex” in Figure 4), RssB undergoes a conformational change that repositions RssB^NTD^, allowing other region(s) of σ^s^ to dock onto RssB^NTD^ in a consolidated interaction (“delivery complex”). At some point, hydrolysis of the high energy acyl phosphate bond will occur, lowering the affinity of RssB for σ^s^ and perhaps signaling substrate hand-off and transition to the elongation mode of ClpXP, at which step ClpX needs to loosen its grip on the substrate and be able to processively move the polypeptide through its pore for unfolding and ultimate transfer to the ClpP chamber. Excessively strong binding of an adaptor at this step may be a disadvantage because it may impede substrate sliding. Thus, it appears likely that at some point, RssB will let go of one of the σ^s^ regions and will eventually completely release it. Building upon our earlier model^34^, we speculate here that the RssB molecule does in fact adopt a more open conformation, but that this conformation requires both phosphorylation and docking of σ^s^. The open form could be achieved by widening of the cleft between RssB^NTD^ and RssB^CTD^, with σ^s^ serving as a spacer that disrupts contacts between the receiver and effector domain, which are generally inhibitory in response regulators^50,51^.

It is worthwhile to note that in some response regulators, phosphorylation-dependent activation is achieved through a second modality of structural rearrangement, dimerization. This is mediated usually, but not always, by the 4-5-5 face as observed in the OmpR and NtrC subfamilies. The potential role of RssB oligomerization in the overall mechanism remains unclear and is not considered in Figure 4. Dimerization has been shown to be part of the mechanism of action of the *Caulobacter crescentus* adaptor, RcdA. RcdA dimers can readily be detected in solution, and critically RcdA and cargo both compete for the same RcdA interface. This competition determines the fate of RcdA itself since RcdA homodimers are degraded by ClpXP, while bound cargo stabilizes RcdA against proteolysis^52,53^.

In contrast, the evidence for RssB dimerization is weaker and cannot be readily formalized into a mechanistic scheme. RssB oligomerization has been detected by bacterial two-hybrid assay in *clpX* and *clpXP*-deleted strains^32^. However, *in vitro*, dimerization and higher order oligomerization has only been detected by chemical crosslinking of full-length RssB^34^ and by gel filtration and X-ray crystallography of isolated RssB^NTD 41^. That said, there are reasons to question whether dimerization is part of the RssB mechanism. First, RssB levels are extremely low in cells^48^ and remain largely constant due to a homeostatic feedback loop^28^. Second, the contacts holding the RssB^NTD^ dimer together are different from those seen in other response regulators^30^ and cannot be ruled out as a crystallization artifact. Third, dimerization has not been seen with full-length RssB via size-exclusion chromatography coupled to multi-angle light scattering detection at concentrations well above physiological, and importantly, phosphorylation does not appreciably alter the oligomerization state of RssB^34^. Altogether, these observations suggest that RssB dimers, if they occur in solution, are transient and may require stabilization by one or more binding partners. Data so far argue against σ^s^ only promoting dimerization^30^. In fact, our own data suggest that σ^s^ does not cause RssB dimerization, as the detected σ^s^:RssB complex in Figure 1A is consistent with a 1:1 stoichiometry. It is possible that under certain conditions, the local concentration of RssB will rise high enough for RssB to dimerize, and that these dimeric species get degraded by ClpXP, rather than recycled. While combined approaches are beginning to unravel many facets of σ^s^ regulation, further structural studies of RssB in complex with its binding partners (IraP, IraM, ClpXP, σ^s^) will be instrumental to satisfactorily dissect the mechanistic basis for this hub of interactions supported by RssB. Our study presents a novel type of phosphorylation-dependent rearrangement that highlights the dynamic role of α5 and of the SHL and points to yet another paradigm for how some members of the large and ancient family of response regulators signal upon phosphorylation.

## METHODS

### RssB Production and Crystallization

RssB and His-tagged RssB were purified by a succession of nickel-affinity chromatography and gel filtration as described by Dorich et al.^34^. For crystallization, we used the non-tagged RssB construct. BeSO_4_ and NaF were premixed for 15 minutes on ice and added to RssB protein to a final concentration of 1 mM beryllofluoride. Protein was dispensed manually in hanging drops on 24-well VDX plates (Hampton Research) and high-quality crystals were obtained with 30 mM ammonium sulfate, 22% w/v PEG 3350. The crystallization cocktail was supplemented with 25% PEG 400 for purposes of cryoprotection. Flash-cooling was achieved by plunging into liquid nitrogen.

### Structure Determination and Refinement

Diffraction data were collected at beamline 24-ID-C (Argonne National Laboratory), integrated and scaled using XDS^54^ and Aimless. The structure was phased using molecular replacement with the homologous domains of PDB ID 6OD1 as search models using the Molecular Replacement Phaser Module in PHENIX^55^. Iterative refinement and model building was performed using PHENIX ^55^ and Coot ^56^. Data collection, refinement statistics are presented in Table 1.

### RssB Phosphorylation Assays

RssB was dialyzed overnight against 20mM Tris-HCl pH8, 150mM NaCl, 15mM MgCl_2_, 10% glycerol, 1mM DTT. Electrophoresis samples contained 1.5μg protein supplemented with 15mM AcP. At indicated timepoints, 10μL samples were withdrawn from the total sample and mixed with 4μL of 4xLDS (0.25M Tris-HCl pH 6.8, 8% SDS, 0.2M DTT, 0.04% bromophenol blue). For electrophoresis, we utilized 10% SuperSep Phos-tag precast gels (Fujifilm Wako), which were run with 1xTris-Tricine buffer (0.1M Tris base, 0.1M Tricine, 0.1% SDS) for 90min at 150V in the cold. Gels were fixed for 10min (50% methanol, 10% acetic acid) and stained with Coomassie Blue for imaging on a gel doc (Bio-Rad). Band intensities were quantified using Image Lab (Bio-Rad) and data plotted using Prism 9 (GraphPad). Experiments were performed in two or more replicates.

### Analytical Size-Exclusion Chromatography

RssB and RpoS were dialyzed overnight against 20mM Tris-HCl pH 8, 250mM NaCl, 10mM MgCl_2_, 2mM β-mercaptoethanol. RssB was phosphorylated by adding 25mM lithium potassium acetyl phosphate (AcP, Sigma) and incubation for 15min on ice. Samples of RssB, σ^s^, RssB+σ^s^, RssB∼P, RssB∼P+σ^s^ were prepared at 15 μM final concentration in a total volume of 100 μl. A Superdex 200 Increase 10/300 column (GE Healthcare) was equilibrated with dialysis buffer (±25mM AcP) and calibrated with molecular weight standards (Bio-Rad) prior to sample loading. The void volume of the column was determined using Dextran Blue. Peak fractions were collected and analyzed by SDS-PAGE on 4-12% Bis-Tris gels (Life Technologies).

### Pull-Down Assays

RssB, RssB variants and His-tagged RpoS were dialyzed overnight against 20mM Tris-HCl pH 8, 250mM NaCl, 10mM MgCl_2_, 15mM imidazole, 2mM β-mercaptoethanol. RssB and RssB variants were treated with 1mM BeF_3_ ^-^ for 15min on ice. Pull-downs were performed in 66uL reactions. His-tagged σ^s^ was added to phosphorylated/unphosphorylated RssB in Micro Bio-Spin Chromatography columns (Bio-Rad) before removing 6μL of the input for SDS-PAGE analysis and adding 40μL of prepared HisPur Ni-NTA resin (Thermo Scientific). Samples were incubated for 60min on a rotating wheel at 4°C before flow through was collected by centrifugation for 1min at 1000 rcf. Resin was washed 3 times with 300μL dialysis buffer and one more time with 40μL of buffer. Phosphorylated samples were washed with dialysis buffer supplemented with 1mM BeF_3_ ^-^. Complexes were eluted by incubation with IMAC-600 (20mM Tris-HCl pH 8, 250mM NaCl, 10mM MgCl_2_, 600mM Imidazole, 2mM β-mercaptoethanol (+/-beryllofluoride) for 5 min at 4°C followed by centrifugation for 2 min at 1000 rcf. Input and elution samples were analyzed by SDS-PAGE on 4-12% Bis-Tris gels (Life Technologies), followed by gel staining with Coomassie Blue and imaging. Band intensities were quantified using Image Lab (Bio-Rad) and plotted using Prism 9 (GraphPad). Experiments were performed in triplicate.

### Intrinsic Tryptophan Fluorescence

Protein samples were dialyzed overnight against 50mM Tris-HCl pH8, 200mM NaCl, 10mM MgCl_2_, 2mM β-mercaptoethanol and diluted to 2μM prior to the experiment. Measurements were recorded in triplicate using a Fluoromax-4 spectrofluorometer (Horiba). 150uL of protein solution +/-1mM beryllofluoride or 6mM NaF were transferred to a quartz cuvette and measurements were taken at 25°C in 1nm increments over a wavelength range of 300-500nm. The excitation wavelength was set to 290nm. Spectra were corrected for background. A NaF control measurement was recorded to ensure that any potential change in signal observed upon incubation with beryllofluoride is not due to excess NaF used in the preparation of the phosphomimic.

### Circular Dichroism

Protein was dialyzed overnight against 20mM Tris-HCl pH8, 50mM NaCl, 10mM MgCl_2_, 1mM TCEP and diluted to 2.5μM prior to the experiment. The assay was performed on a Jasco J-815 CD spectrometer. Measurements were performed at 25°C in a 2mm pathlength cuvette over a wavelength range of 200-260nm with a speed of 50nm/min. Samples were measured in triplicate with 3 acquisitions per sample and corrected for background.

## ACKNOWLEDGEMENTS

Research in the laboratory of A.M.D. was supported by the National Institutes of Health (R01GM121975; R35GM144124) and a Salomon Research Award from Brown University. This work used NE-CAT beamlines (GM124165) at the Advanced Photon SOURCE (DE-AC02-06CH11357) and the Proteomics Facility at Brown University. We thank staff at NECAT 24-ID for excellent beamline support and Ms. V. Frankovich for technical assistance during the early stages of the project. We thank Dr. S. Gottesman and Dr. Sue Wickner for critical reading of the manuscript.

## CONFLICT OF INTEREST

The authors declare no conflict of interest.

## AUTHOR CONTRIBUTION

C.B. collected diffraction data and carried out protein refinement and *in vitro* characterization. J.S. purified protein and obtained crystals. M.F. developed RssB phosphorylation assays. A.M.D designed the overall study, oversaw data processing and analysis, acquired funding and wrote the initial draft, which was edited and approved by all authors.

## FIGURE LEGENDS

**Movie 1**. The movie shows the structure of IraD-bound RssB^D58P^ morphing into the structure of beryllofluoride-bound RssB reported here. RssB domains are colored as in Figure 2A and IraD is green.

## Notes

### Competing Interest Statement

The authors have declared no competing interest.

